# Differentiating the Roles of UL16, UL21 and Us3 in the Nuclear Egress of Herpes Simplex Virus Capsids

**DOI:** 10.1101/2020.02.14.949750

**Authors:** Jie Gao, Renée L. Finnen, Maxwell R. Sherry, Valerie Le Sage, Bruce W. Banfield

## Abstract

Previous studies from our laboratory established that pUL16 and pUL21 are required for efficient nuclear egress of herpes simplex type 2 (HSV-2) capsids. To better understand the role of these proteins in nuclear egress, we wished to establish whether nuclear egress complex (NEC) localization and/or function was altered in the absence of either pUL16 or pUL21. We used antiserum raised against HSV-2 NEC components pUL31 and pUL34 to examine NEC localization by immunofluorescence microscopy. NEC localization in cells infected with pUL16 deficient viruses was indistinguishable from that observed in cells infected with wild type viruses. By contrast, NEC localization was found to be aberrant in cells infected with pUL21 deficient virus and, instead, showed some similarity to the aberrant NEC localization pattern observed in cells infected with pUs3 deficient virus. These results indicated that pUL16 plays a role in nuclear egress that is distinct from that of pUL21 and pUs3. Higher resolution examination of nuclear envelope ultrastructure in cells infected with pUL21 deficient viruses by transmission electron microscopy showed different types of nuclear envelope perturbations, including some that were not observed in cells infected with pUs3 deficient virus. The formation of the nuclear envelope perturbations observed in pUL21 deficient virus infections was found to be dependent on a functional NEC, revealing a novel role for pUL21 in regulating NEC activity. The results of comparisons of nuclear envelope ultrastructure in cells infected with viruses lacking pUs3, pUL16 or both pUs3 and pUL16 were consistent with a role for pUL16 upstream of primary capsid envelopment and shed new light on how pUs3 functions in nuclear egress.

**Author summary:** The membrane deformation activity of the herpesvirus nuclear egress complex (NEC), allows viral capsids to transit from their site of assembly in the nucleus through both nuclear membranes into the cytoplasm. The timing, extent and directionality of NEC activity must be precisely controlled during viral infection, yet our knowledge of how NEC activity is controlled is incomplete. To determine how pUL16 and pUL21, two viral proteins required for nuclear egress of herpes simplex virus type 2 (HSV-2) capsids, function to promote nuclear egress, we examined how the lack of each protein impacted NEC localization. These analyses revealed a function of pUL16 in nuclear egress that is distinct from that of pUL21, uncovered a novel role for pUL21 in regulating NEC activity and shed new light on how a viral kinase, pUs3, regulates nuclear egress. Nuclear egress of viral capsids is a common feature of the replicative cycle of all herpesviruses. A complete understanding of all aspects of nuclear egress, including how viral NEC activity is controlled, may yield strategies to disrupt this process that could be applied to the development of herpes-specific antiviral drugs.

## Introduction

Herpes virions are complex structures comprised of a large DNA genome encapsulated in an icosahedral capsid surrounded by a proteinaceous layer of tegument, which in turn is surrounded by a lipid envelope studded with numerous membrane glycoproteins. Befitting this structural complexity, the morphogenesis of these particles is equally complex and is not fully understood. The virion assembly pathway begins in the infected cell nucleus where newly replicated viral genomes are packaged into preformed procapsids. DNA-containing capsids are then preferentially selected for further maturation and acquire a primary envelope by budding into the inner nuclear membrane (INM) resulting in the formation of a primary enveloped virion (PEV) residing in the perinuclear space. These PEVs then de-envelop at the outer nuclear membrane (ONM) releasing a capsid into the cytoplasm that is subsequently enveloped at a post-Golgi compartment and the mature virion is secreted from the cell (1).

The present study concerns the nuclear egress of capsids from the nucleoplasm to the cytoplasm. Key viral factors required for this process are the components of the nuclear egress complex (NEC). Components of the NEC, the orthologs of the herpes simplex virus (HSV) proteins pUL31 and pUL34, localize predominantly to the INM in virally infected cells (2, 3). Studies from several laboratories have described the molecular structure of NEC complexes derived from a number of herpesviruses and these structures have provided important insight into the molecular mechanisms by which the NEC promotes capsid envelopment at the INM (4–9). Deletion of UL31 or UL34 from HSV-1 results in a 4-log reduction in virus replication in most cell types, and the accumulation of capsids in the nuclei of infected cells (10, 11). pUL34 is a type II membrane protein with a C-terminal transmembrane domain. The N-terminus of pUL34 is localized on the cytoplasmic or nucleoplasmic face of the endoplasmic reticulum and INM, respectively. pUL31, a soluble nucleoplasmic phosphoprotein, can interact with the N-terminus of pUL34 and is recruited to the INM in the presence of pUL34. Expression of pUL31 and pUL34 in the absence of other viral proteins leads to vesiculation of the INM at the nuclear periphery (12) and the interaction of purified pUL31 and pUL34 with synthetic membranes in vitro results in the induction of membrane vesiculation (13). Artificial targeting of purified pUL31 to membranes in the absence of pUL34 also leads to membrane vesiculation in vitro, suggesting that a key function of pUL34 is to recruit pUL31 to the membrane surface (14). In infected cells, however, pUL31 and pUL34 are distributed evenly throughout the INM and vesiculation of the INM is not observed (2, 12). These findings suggest that the INM vesiculation activity of the NEC is negatively regulated by a protein(s) found in infected cells.

One protein that is capable of regulating NEC activity is the viral serine/threonine kinase pUs3. In cells infected with pUs3 mutants of pseudorabies virus (PRV), HSV-1, equine herpes virus 1 (EHV-1) and Marek’s disease virus (MDV) PEVs accumulate in herniations of the INM (3, 15–17). HSV-1 pUs3 phosphorylates pUL31 at a number of serine residues in its N-terminus and it has been proposed that these modifications are required for efficient de-envelopment of PEVs at the ONM, because mutation of these pUL31 serine residues to alanine recapitulates the pUs3 mutant phenotype (18). Moreover, mutation of the serine residues in the N-terminus of HSV-1 pUL31 to glutamic acid, to mimic the hyper-phosphorylated pUL31 state, resulted in a failure of capsids to be efficiently enveloped at the INM (18). Taken together, these findings suggest that a hypo-phosphorylated form of pUL31 is required for NEC mediated envelopment of capsids and that pUs3 may act to restrict NEC mediated INM vesiculation by phosphorylating pUL31.

Previous studies from our laboratory demonstrated that HSV-2 strain 186 lacking either pUL21 or pUL16 showed deficiencies in the nuclear egress of capsids (19–21). As pUL16 and pUL21 can form a complex (22), we hypothesized that this complex was required for efficient nuclear egress of capsids (19). To define the roles of pUL16 and pUL21 in nuclear egress and to determine whether these proteins function individually or as a complex in nuclear egress, we analyzed the localization and function of the NEC in cells infected with multiple HSV strains deficient in either pUL16 or pUL21. Our analyses revealed that pUL16 mutants were phenotypically distinct from pUL21 mutants in terms of NEC localization in infected cells and indicated that pUL16 functions upstream of pUL21 and pUs3 in nuclear egress. Our analyses also revealed a novel role for pUL21 in regulating the activity of the NEC. Based on the results of further phenotypic analyses of viruses lacking both pUL16 and pUs3, we also propose an new interpretation of pUs3 function in nuclear egress.

## Results

### The NEC is mislocalized in cells infected with HSV strains lacking pUL21 or pUs3

One possible explanation for the capsid nuclear egress deficiencies seen with HSV-2 pUL21 and pUL16 mutants is that these proteins, individually or as a complex, are required for the appropriate localization and/or function of the viral nuclear egress complex (NEC), comprised of pUL31 and pUL34. To enable investigation of HSV-2 NEC localization and function, antisera reactive against HSV-2 pUL31 and pUL34 were produced in rat and chicken, respectively. The antisera produced were highly specific in both Western blot analyses (Fig 1A) and in indirect immunofluorescence microscopy experiments (Fig 1B) and were reactive against pUL31 and pUL34 orthologs from both HSV-2 and HSV-1 (see below).

**Fig 1.**
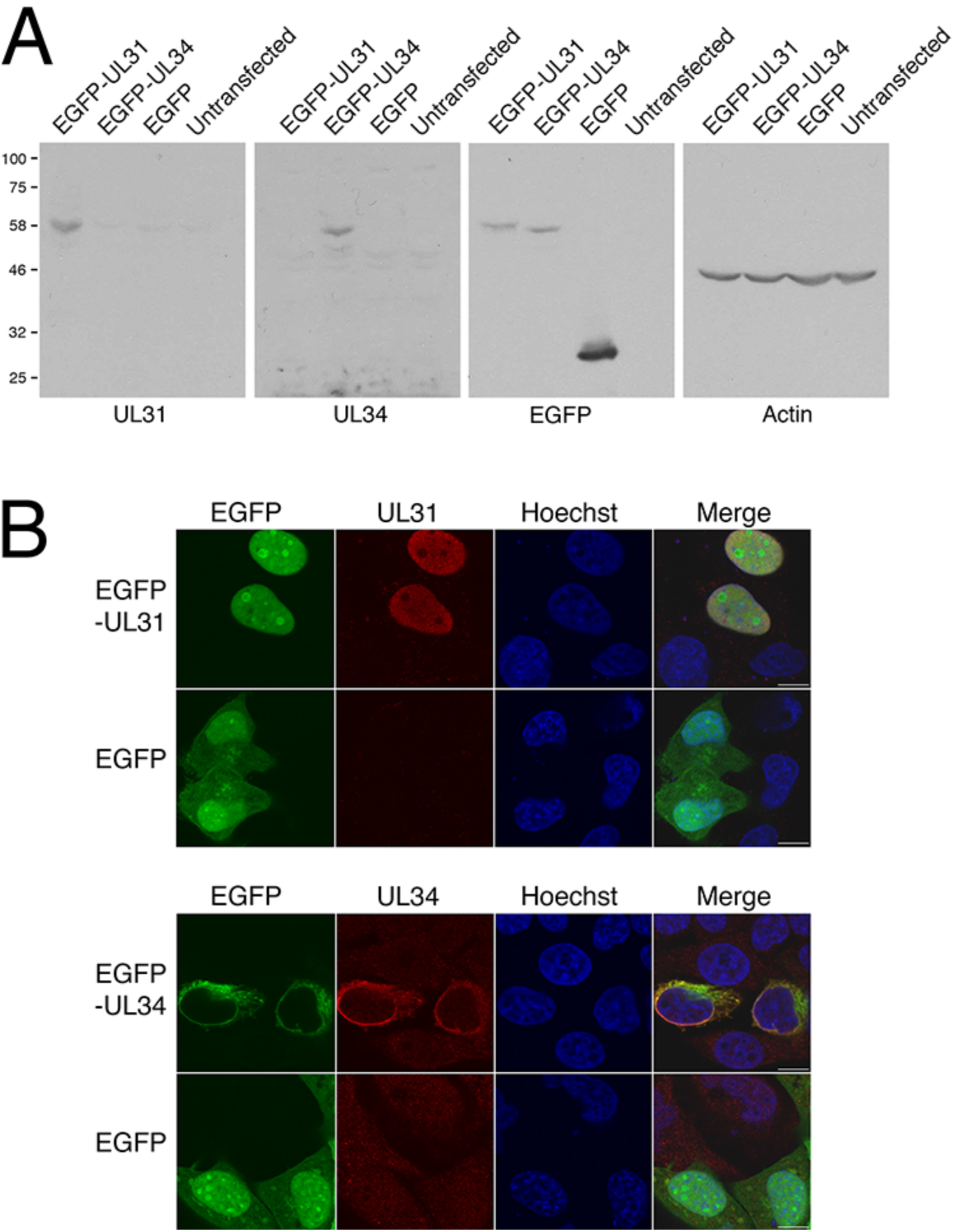
Characterization of antisera generated against HSV-2 pUL31 and HSV-2 pUL34. **(A)** Western blot analysis. Equal volumes of whole cell protein extracts prepared at 24 hours post transfection from 293T cells transfected with plasmids encoding the proteins indicated at the top of each panel, or from untransfected 293T cells, were electrophoresed through 8% polyacrylamide gels and transferred to PVDF membranes. Membranes were probed with antisera indicated at the bottom of each panel. Molecular weight markers in kDa are indicated on the left side of the panels. (**B)** Indirect immunofluorescence microscopy. HeLa cells were transfected with plasmids encoding the proteins indicated at the left side of each panel then fixed at 18 hours post transfection and incubated with either pUL31 antisera (top panels) or pUL34 antisera (bottom panels) followed by incubation with the appropriate Alexa Fluor 568-conjugated secondary antibody. Nuclei were stained with Hoechst 33342. Scale bars are 10 μm.

Using these antisera we investigated the localization of the NEC in cells infected with pUL21, pUL16 and pUs3 deficient viruses derived from HSV-2 strain 186 (designated hereafter as Δ21, Δ16 and ΔUs3). Whereas NEC localization was uniform and smooth at the nuclear rim of cells infected with WT and Δ16 strains, the NEC was mislocalized in cells infected with Δ21 and ΔUs3 (Fig 2). In the case of Δ21 infected cells, pUL31 and pUL34 still co-localized at the nuclear rim, however the localization pattern was uneven with puncta of varying size distributed irregularly on both the nucleoplasmic and cytoplasmic sides of the nuclear envelope. The pattern of NEC localization in ΔUs3 infected cells appeared distinct from that observed in Δ21 infected cells, with a more uniform, regularly spaced arrangement of puncta along the nuclear rim. A similar pattern of NEC localization has been observed in cells infected with Us3 deficient strains of several alphaherpesviruses (3, 15–17). The differences in NEC localization observed between Δ16, Δ21 and ΔUs3 HSV-2 strains suggests that pUL16, pUL21 and pUs3 play roles that are distinct from one another in the nuclear egress of HSV-2. Thus, our earlier hypothesis that pUL16 and pUL21 functioned as a complex to promote nuclear egress is likely incorrect (19).

**Fig 2.**
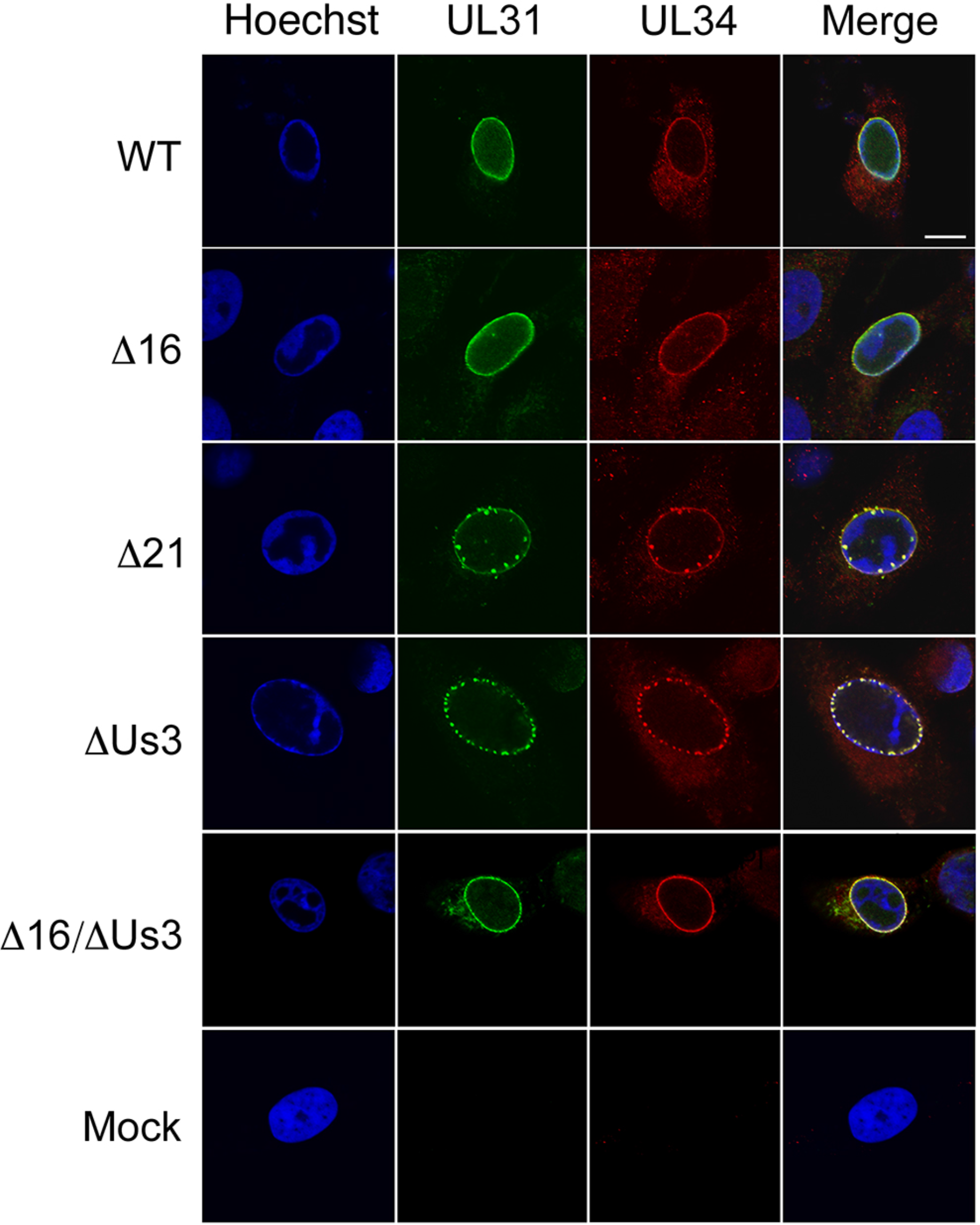
Localization of the NEC in cells infected with WT and mutant HSV-2 strains. Vero cells were infected with the indicated viruses or mock-infected. At 8 hpi, cells were fixed and stained with pUL31 and pUL34 antisera and Alexa Fluor 488-conjugated and Alexa Fluor 568-conjugated secondary antibodies, respectively. Nuclei were stained with Hoechst 33342. Scale bar is 10 μm.

To investigate whether the striking mislocalization of the NEC observed in cells infected with Δ21 was a phenotype common to other strains of HSV-2 and HSV-1, we examined the localization of the NEC in cells infected with two additional strains of HSV-2 (HG52 and SD90e) and two strains of HSV-1 (F and KOS) alongside their corresponding pUL21 mutants (23). Cells infected with pUL21 deficient viruses, regardless of background strain, showed similar aberrant NEC localization in comparison to cells infected with parental viruses (Fig 3). These findings indicate that NEC mislocalization is a conserved property of pUL21 mutants across HSV species and strains.

**Fig 3.**
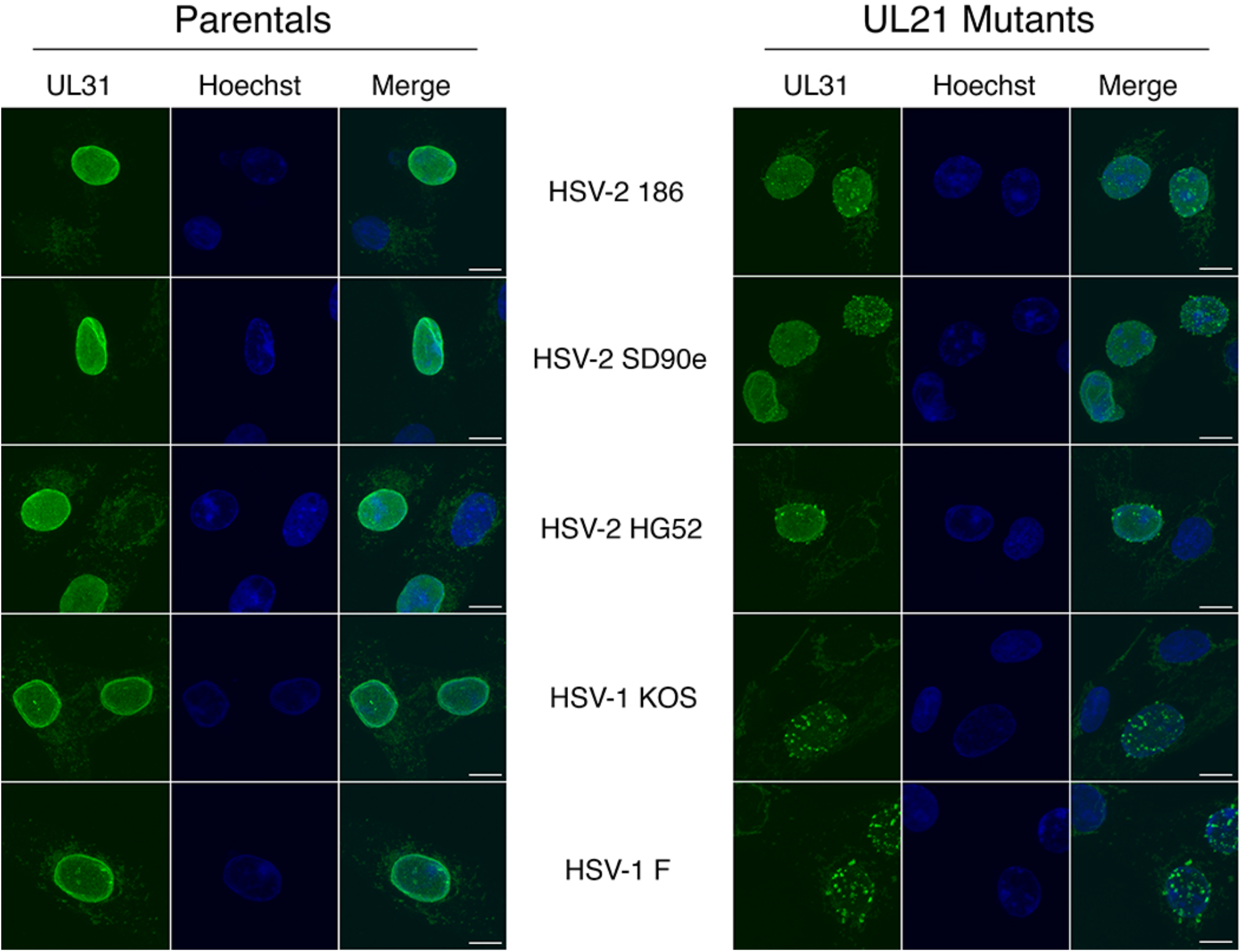
Localization of the NEC is aberrant in cells infected with virus lacking pUL21 regardless of background strain. Vero cells were infected with the indicated parental strains and their corresponding mutants lacking pUL21. At 8 hpi, cells were fixed and stained with pUL31 antisera and Alexa Fluor 488-conjugated secondary antibodies. Nuclei were stained with Hoechst 33342. Z-series projection images, generated with FV10-ASW version 04.01 software, are shown. Scale bars are 10 μm.

To ensure that this NEC mislocalization was due to loss of pUL21 and not due to unintended mutations introduced during strain construction, HaCaT cells stably expressing HSV-2 pUL21, HaCaT21, were isolated and infected with WT and pUL21 mutant HSV strains derived from HSV-2 (Δ21) and HSV-1 (FFS62/34). Whereas parental HaCaT cells infected with pUL21 mutants displayed NEC mislocalization similar to that observed in infected Vero cells (compare Fig 4A with Fig 3), HaCaT21 cells infected with pUL21 mutants displayed smooth NEC localization (Fig 4A). The proportion of infected cells displaying smooth versus perturbed NEC localization was quantified in two independent experiments (Fig 4B). Whereas expression of HSV-2 pUL21 in HaCaT cells had no effect on the localization of the NEC in WT virus infected cells, it restored the normal, smooth, localization of the NEC in pUL21 mutant infected cells from 37.5% to 84.5% in the case of Δ21 and from 7.5% to 41% in FFS62/34 infected cells. The superior complementation observed with the HSV-2 versus the HSV-1 UL21 mutant may be due to the expression of a HSV-2 pUL21 ortholog in HaCaT21 cells rather than a HSV-1 derived pUL21. Taken together, these data confirm that the perturbation in NEC localization observed in cells infected with pUL21 mutant strains was due to the loss of pUL21.

**Fig 4.**
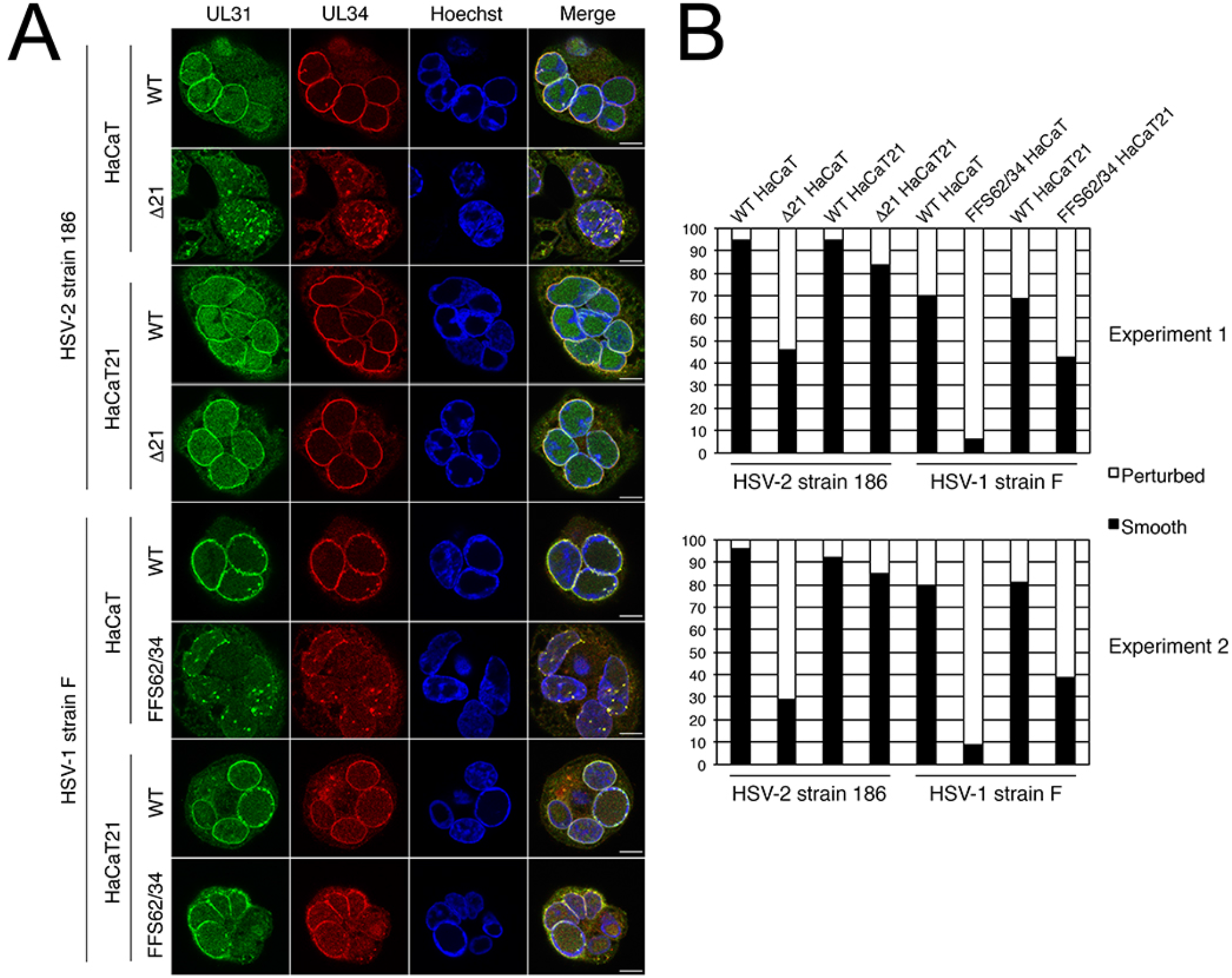
HaCaT21 cells complement the aberrant NEC localization phenotype of viruses lacking pUL21. **(A)** HaCaT or HaCaT21 cells were infected with the indicated virus strains. At 18 hpi cells were fixed and stained with pUL31 antisera and pUL34 antisera and Alexa Fluor 488-conjugated and Alexa Fluor 568-conjugated secondary antibodies, respectively. Nuclei were stained with Hoechst 33342. Scale bars are 10 μm. **(B)** To quantify NEC localization (perturbed versus smooth), one hundred infected cells were scored for the appearance of pUL34 around the nuclear rim. Two independent experiments were scored.

### Cells infected with HSV strains lacking pUL21 display a variety of nuclear envelope perturbations

As the NEC is anchored within the INM by its pUL34 component, the NEC labeled irregular puncta in UL21 mutant infected cells that appeared to protrude into and out of the nucleus likely represented perturbations of the nuclear envelope. To examine this at higher resolution, transmission electron microscopy (TEM) analyses were carried out on Vero cells infected with three strains of HSV-2 (186, HG52 and SD90e) and two strains of HSV-1 (F and KOS) alongside their corresponding pUL21 mutants. The appearance of the nuclear envelope in cells infected with WT strains were similar to each other, characterized by a smooth appearance occasionally punctuated by small, localized perturbations associated with egressing capsids (Fig 5, arrowheads). By contrast, cells infected with UL21 mutant strains displayed a variety of nuclear envelope perturbations (Fig 6). Extravagations of the nuclear envelope into the cytoplasm were frequently observed and often were comprised of multiple layers of the nuclear envelope arranged in a concentric fashion (Fig 6, white arrowheads), occasionally surrounding cytoplasmic contents that included virus capsids (Fig 6, white arrows). Invaginations of the nuclear envelope into the nucleoplasm, resembling type II nucleoplasmic reticulum were also observed (24) (Fig 6, black arrowheads) as were invaginations of the INM (Fig 6, black arrows), resembling type I nucleoplasmic reticulum that occasionally contained PEVs (24) (Fig 6, white asterisks). These findings are quantified in Figure 7. These analyses demonstrated that perturbations of the nuclear envelope were more readily observed in cells infected with pUL21 deficient viruses, regardless of background strain and explained the perturbations in NEC localization observed in Figures 2, 3 and 4.

**Fig 5.**
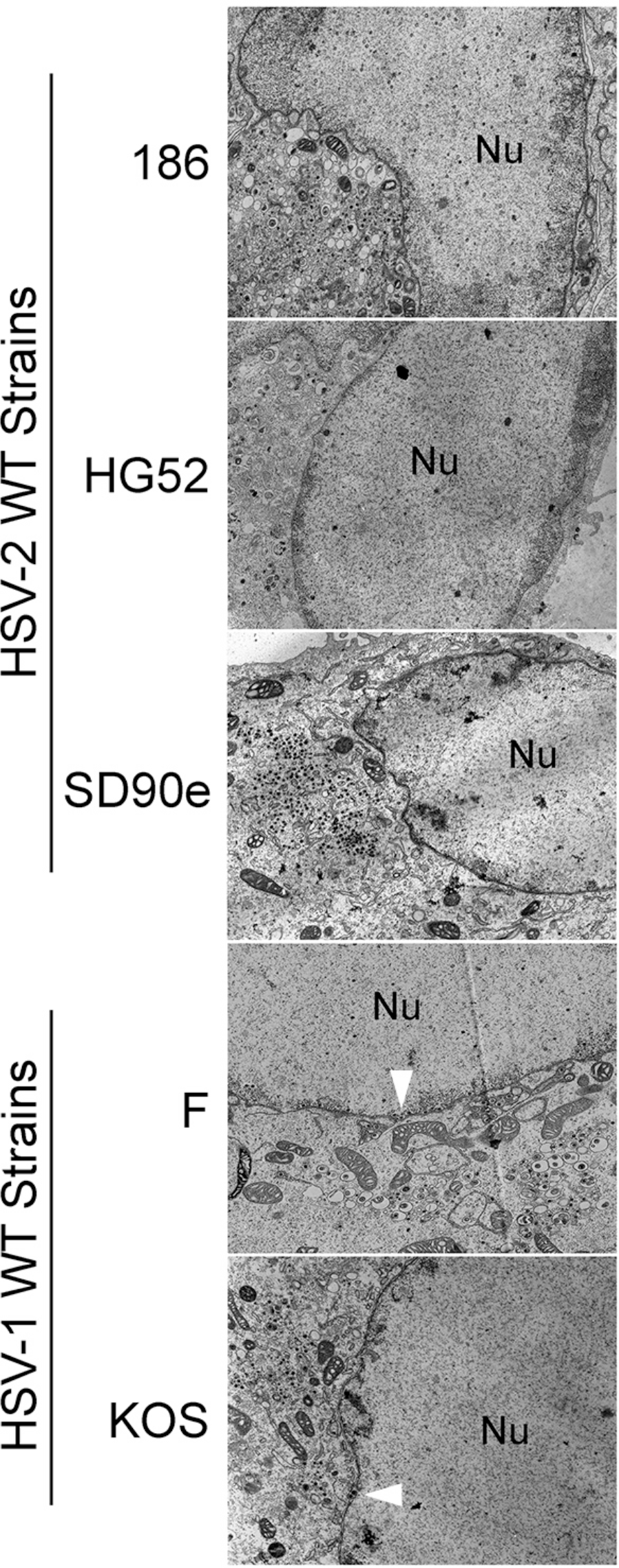
Ultrastructural analysis of cells infected with WT HSV strains. Vero cells were infected with the indicated viruses at an MOI of 3. At 16 hpi, cells were fixed and processed for TEM as described in Materials and Methods. Representative images are shown. Arrowheads indicate small localized nuclear membrane perturbations associated with egressing capsids that were occasionally observed in WT infected cells.

**Fig 6.**
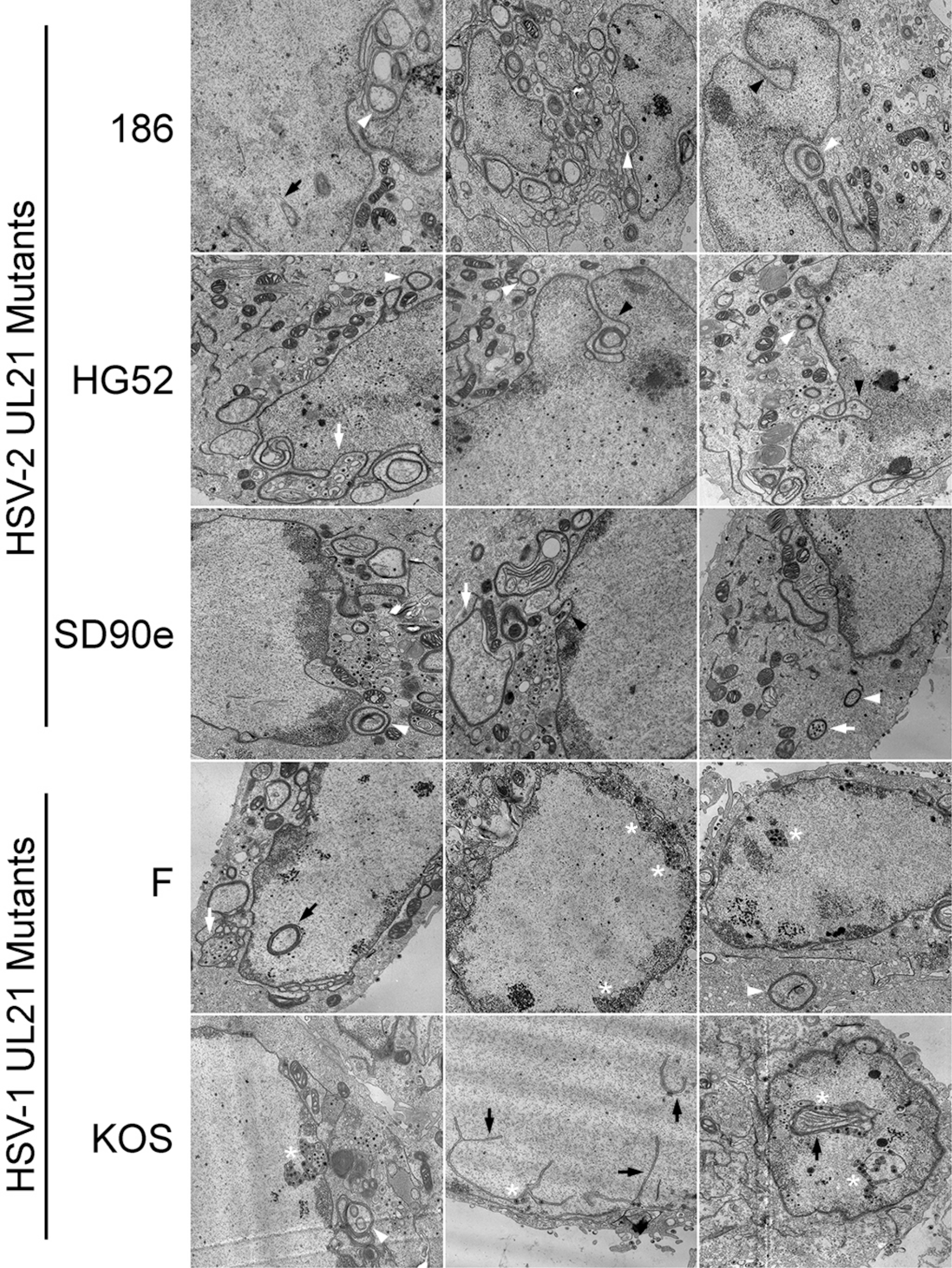
Ultrastructural analysis of cells infected with pUL21 deficient HSV strains. Vero cells were infected with the indicated viruses at an MOI of 3. At 16 hpi, cells were fixed and processed for TEM as described in Materials and Methods. A montage of images depicting the variety of nuclear perturbations that were readily observed in cells infected with pUL21 deficient viruses is shown. White arrowheads indicate extravagations of the nuclear envelope into the cytoplasm, white arrows indicate extravagations surrounding cytoplasmic capsids, black arrowheads indicate invaginations of both membranes of the nuclear envelope into the nucleoplasm, black arrows indicate invaginations of the INM and white asterisks indicate invaginations of the INM containing PEVs.

**Fig 7.**
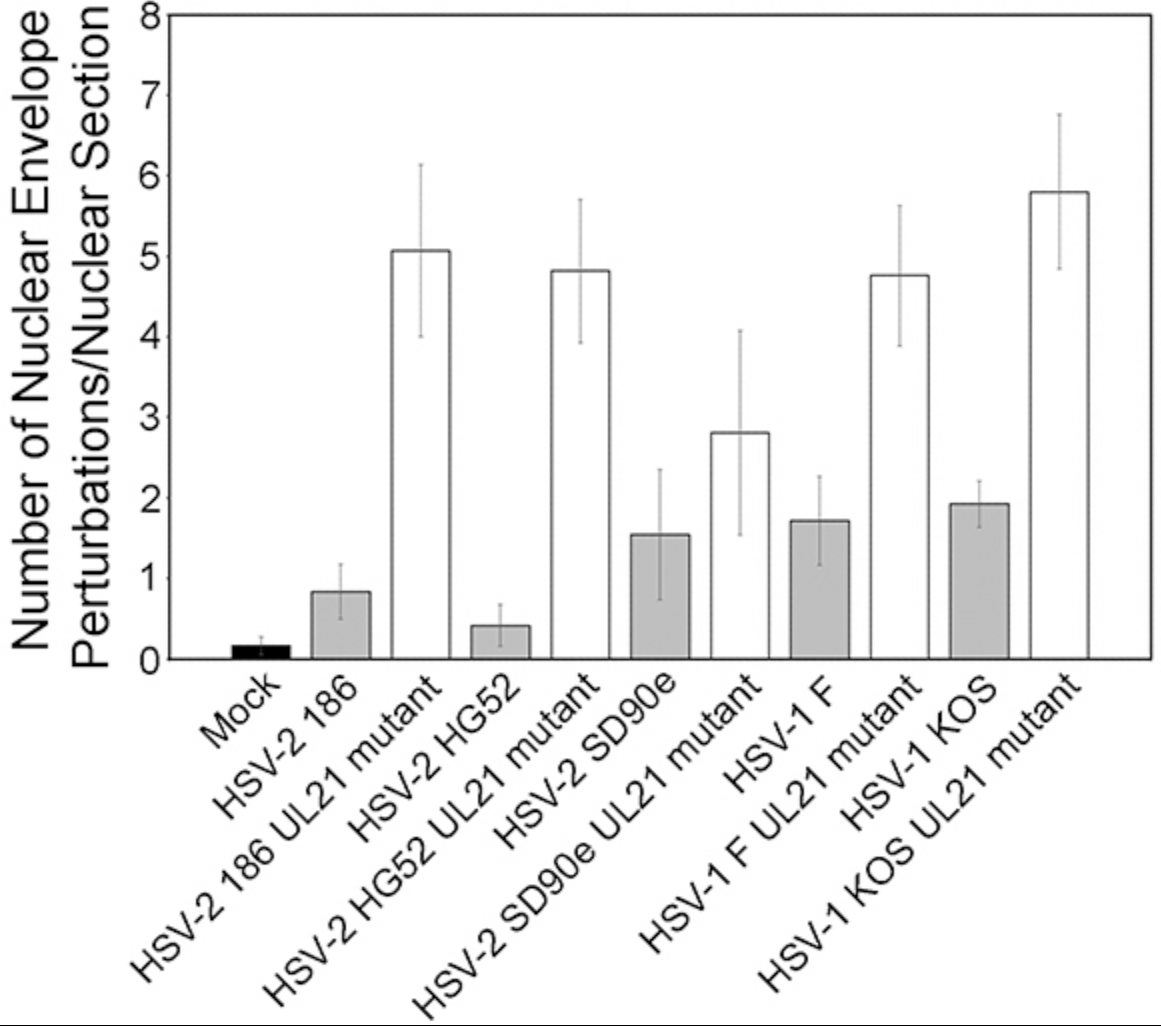
Quantitation of ultrastructural analysis of cells infected with WT or pUL21 deficient HSV. The total number of nuclear envelope perturbations observed in nuclear sections was scored in mock-infected Vero cells and in Vero cells infected with the indicated viruses. More nuclear envelope perturbations were observed in cells infected with pUL21 deficient HSV in comparison to cells infected with the corresponding WT HSV, regardless of background strain. The number of nuclear sections scored were: Mock (n=12); WT 186 (n=12); pUL21 deficient 186 (n=13); WT HG52 (n=12); pUL21 deficient HG52 (n=11); WT SD90e (n=11); pUL21 deficient SD90e (n=16); WT F (n=14); pUL21 deficient F (n=17); WT KOS (n=13); and pUL21 deficient KOS (n=15). Error bars are standard error of the mean.

### The nuclear envelope perturbations associated with Δ21 infection require pUL31

To investigate whether the nuclear envelope perturbation phenotype observed in the absence of pUL21 required a functional NEC, we measured the effect of pUL31 knockdown on nuclear envelope perturbations. Production of pUL31 was specifically knocked down by DsiRNA 19 directed against the UL31 transcript (Figs 8A and B). Whereas knockdown of pUL31 had no effect on the localization of pUL34 at the nuclear rim in WT infected cells, knockdown of pUL31 had a dramatic effect on pUL34 localization in Δ21 infected cells. This effect is most clearly demonstrated in Figure 8C. The cell producing pUL31 (arrowhead) displayed perturbed pUL34 localization whereas the cell knocked down for pUL31 (arrow) displayed smooth pUL34 localization. The proportion of infected cells displaying smooth versus perturbed pUL34 localization was quantified in two independent experiments (Fig 8D). In WT virus infected cells, smooth localization of pUL34 predominated (>90%) in the presence of either DsiRNA 19 or non-silencing control DsiRNA. In Δ21 infected cells, smooth localization of pUL34 was observed at a level comparable to that in WT infected cells in the presence of DsiRNA 19 (92%) but not in the presence of non-silencing control DsiRNA (45.5%). These data are consistent with a requirement for a functional NEC to produce the nuclear envelope perturbations observed in Δ21 infected cells.

**Fig 8.**
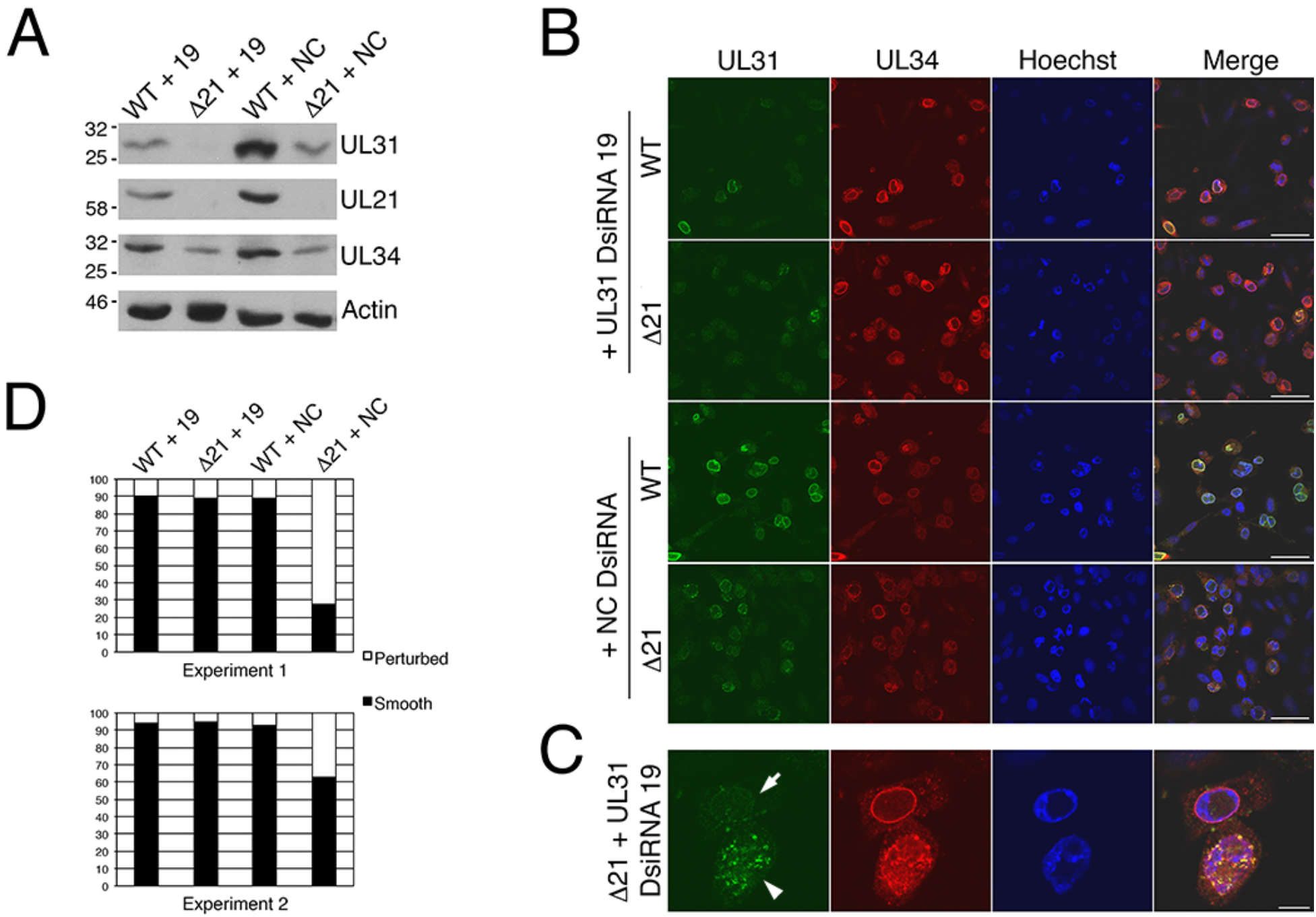
The nuclear membrane perturbation phenotype of viruses lacking pUL21 requires pUL31. Vero cells were transfected with DsiRNAs specific for UL31 (DsiRNA 19) or non-silencing control DsiRNAs (NC) and infected with the indicated viruses at an MOI of 0.5 at 24 hours post transfection. (**A)** At 18 hpi, whole cell extracts were prepared and electrophoresed through 10% polyacrylamide gels and transferred to PVDF membranes. Membranes were probed with antisera indicated on the right side of each panel. Molecular weight markers in kDa are indicated on the left side of each panel. (**B)** At 18 hpi, cells were fixed and stained with pUL31 antisera and pUL34 antisera and Alexa Fluor 488-conjugated and Alexa Fluor 568-conjugated secondary antibodies, respectively. Nuclei were stained with Hoechst 33342. Scale bars are 50 μm. **(C)** Enlarged images of cells transfected with DsiRNA 19 and infected with Δ21 are shown. The arrow indicates a cell in which the production of pUL31 was knocked down and the arrowhead indicates a cell that produced pUL31. Scale bar is 10 μm. **(D)** Quantitation of the effect of UL31 knockdown on the distribution of pUL34 around the nuclear rim. One hundred pUL34 positive cells were scored for the appearance of pUL34 around the nuclear rim (perturbed versus smooth). Two independent experiments were scored.

### HSV-2 pUL16 functions upstream of pUs3 in nuclear egress

As stated above, the smooth NEC localization pattern observed in Δ16 infected cells (Fig 2) supports the notion that pUL16 functions upstream of pUs3 and pUL21 in capsid nuclear egress. A smooth NEC localization pattern was also observed in cells infected with other strains of HSV-2 and HSV-1 lacking pUL16 (20) (Fig 9) confirming that this NEC localization phenotype is conserved amongst HSV species and strains. To further test the idea that pUL16 functions upstream of pUs3, we analyzed the phenotype of two independently isolated HSV-2 strain 186 viruses carrying deficiencies in both proteins (Δ16/ΔUs3). We predicted that if pUL16 was functioning upstream of pUs3 in capsid nuclear egress, the NEC localization in cells infected with the Δ16/ΔUs3 mutants would resemble that of Δ16. Western blot analysis of cells infected with Δ16/ΔUs3 viruses alongside cells infected with WT, Δ16, or ΔUs3 viruses demonstrated the absence of both pUL16 and pUs3 in the double mutants (Fig 10). The localization of the NEC in cells infected with Δ16/ΔUs3 was mostly smooth (Fig 2), similar to what was observed in WT and Δ16 infected cells (Figs 2 and 9). However, closer examination of the nuclear envelope by TEM revealed herniations of the INM that were rarely seen in Δ16 infected cells (Fig 11). These herniations resembled the INM structures formed in ΔUs3 infected cells (Fig 11A, B) except that in Δ16/ΔUs3 infected cells they lacked PEVs (Fig 11C,D). Collectively, these findings are consistent with the idea that that pUL16 functions upstream of capsid primary envelopment. Moreover, the data suggest that it is the lack of pUs3 that is the cause of INM herniation rather than the accumulation of PEVs in the perinuclear space.

**Fig 9.**
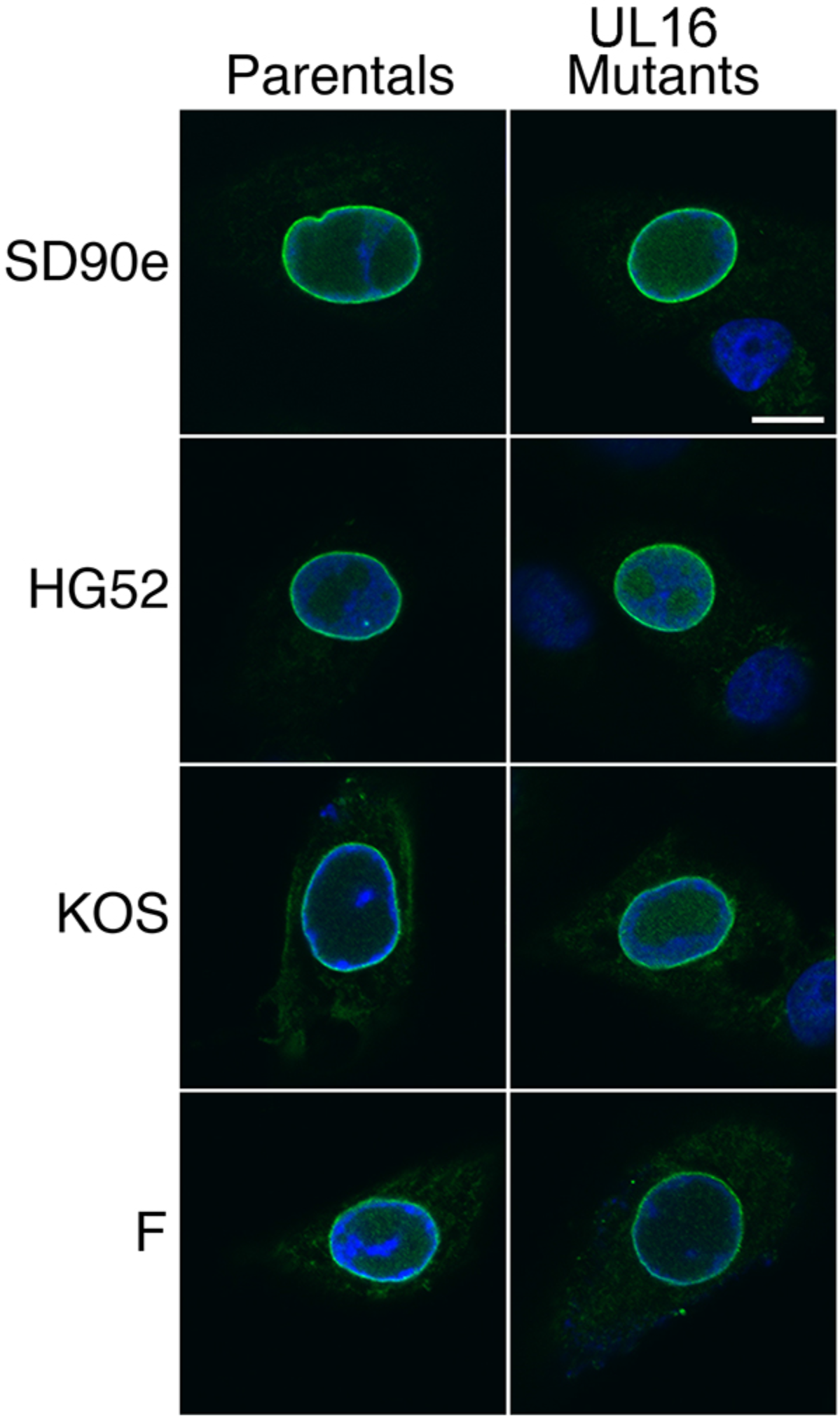
Localization of the NEC is smooth in cells infected with virus lacking pUL16 regardless of background strain. Vero cells were infected with the indicated parental strains and their corresponding mutants lacking pUL16. At 8 hpi, cells were fixed and stained with pUL31 antisera and Alexa Fluor 488-conjugated secondary antibodies. Nuclei were stained with Hoechst 33342 reagent. Scale bar is 10 μm.

**Fig 10.**
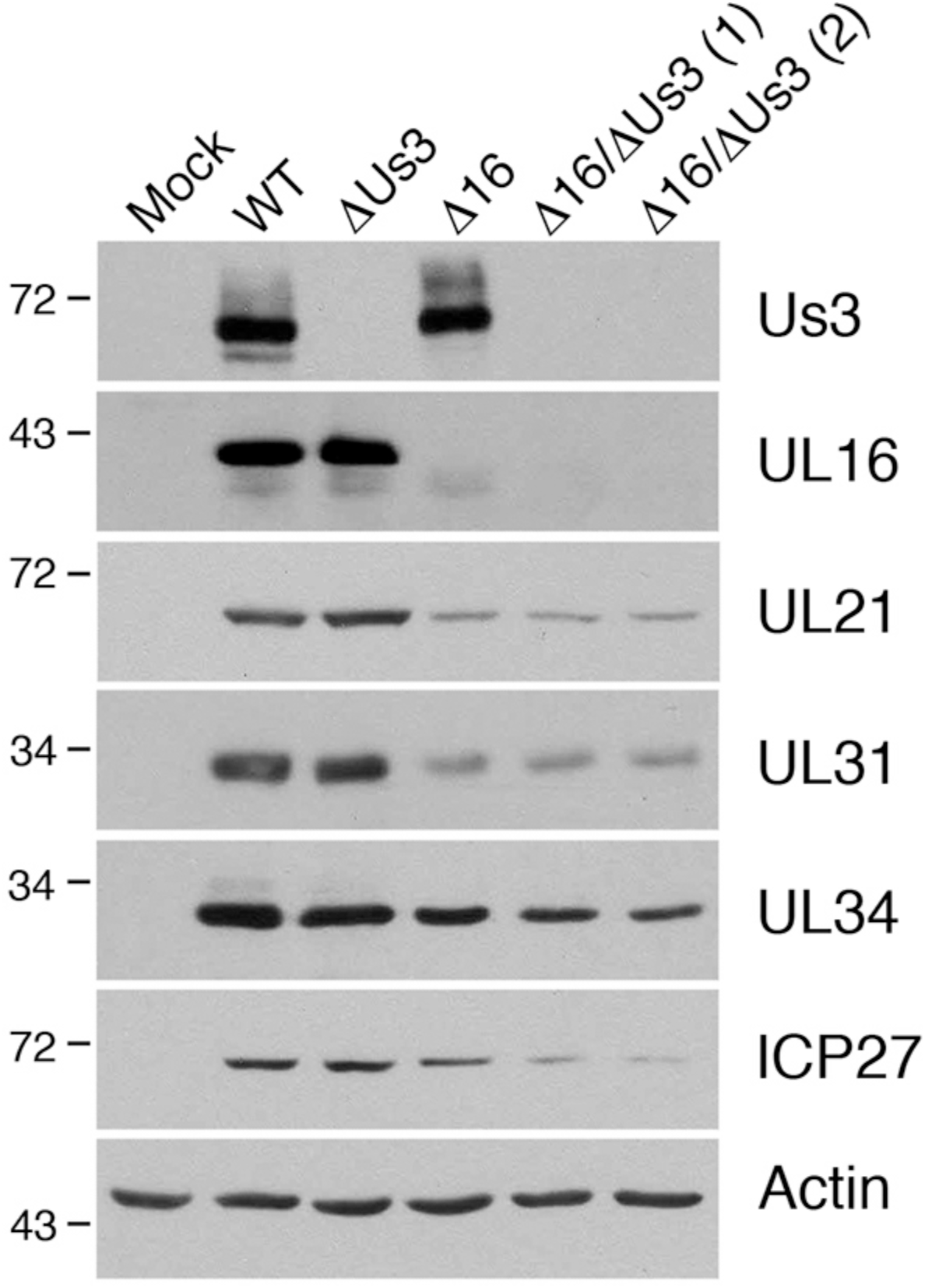
Characterization of Δ16/ΔUs3 mutant virus strains. Vero cells were infected with the indicated viruses at an MOI of 0.1. At 18 hpi, whole cell extracts were prepared and electrophoresed through 10% polyacrylamide gels and transferred to PVDF membranes. Membranes were probed with antisera indicated on the right side of each panel. Molecular weight markers in kDa are indicated on the left side of each panel.

**Fig 11.**
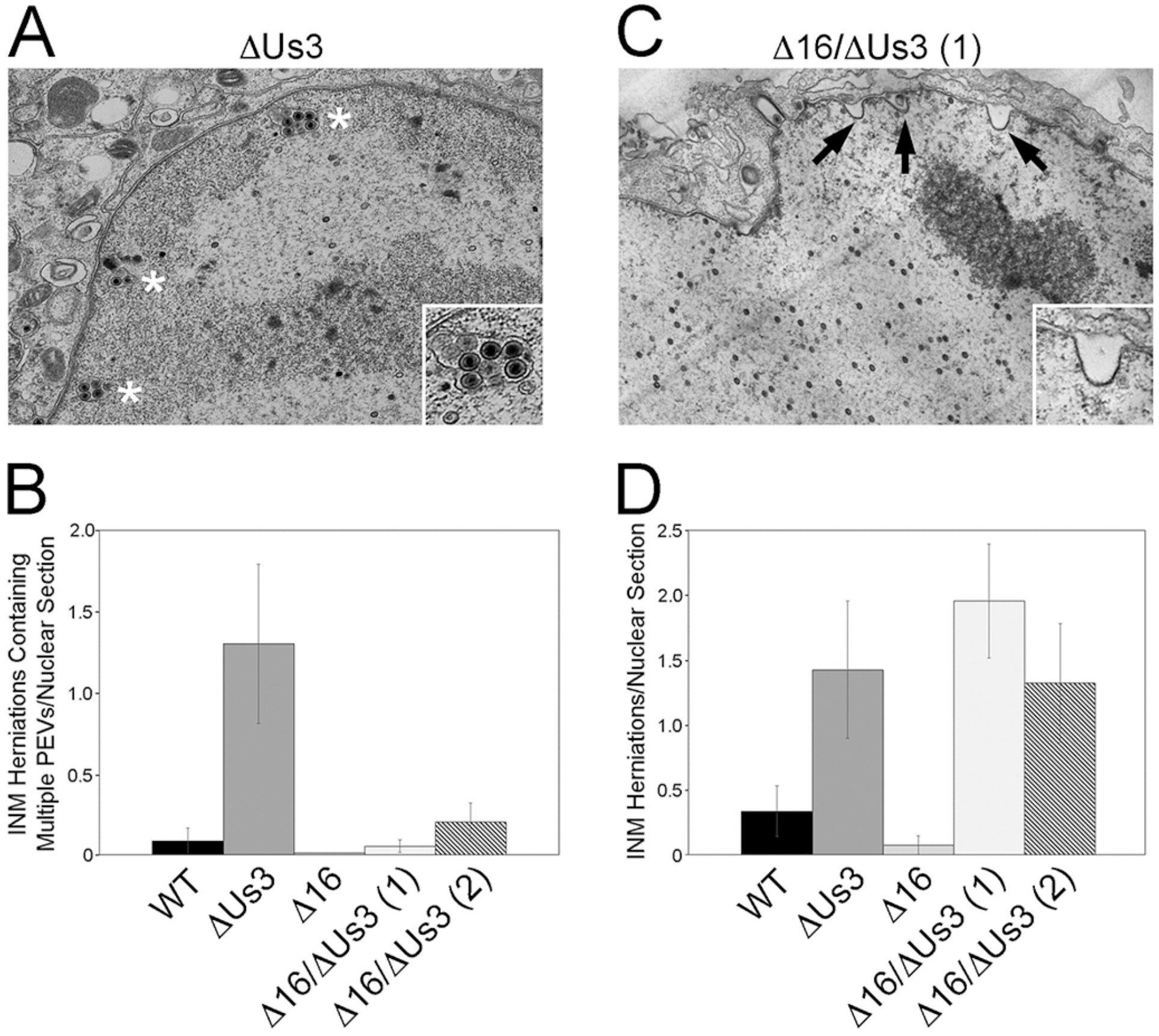
Ultrastructural analysis of cells infected with pUs3 and pUL16 deficient HSV-2 strains. Vero cells were infected with the indicated viruses at an MOI of 3. At 16 hpi, cells were fixed and processed for TEM as described in Materials and Methods. (**A)** Representative image of INM herniations containing PEVs, indicated with asterisks, observed in cells infected with virus deficient in pUs3. An enlarged example of an INM herniation containing PEVs is shown in the inset image. **(B)** Quantitation of INM herniations containing PEVs in infected cells. INM herniations containing PEVs were more frequently observed in cells infected with pUs3 deficient virus than in cells infected with WT virus or pUL16 deficient viruses. The number of nuclear sections scored were: WT 186 (n=12); ΔUs3 (n=28); Δ16 (n=14); Δ16/ΔUs3 isolate 1 (n=38) and Δ16/ΔUs3 isolate 2 (n=10). Error bars are standard error of the mean. **(C)** Representative image of INM herniations lacking PEVs, indicated with black arrows, observed in cells infected with virus deficient in both pUs3 and pUL16. An enlarged example of an INM herniation lacking PEVs is shown in the inset image. **(D)** Quantitation of INM herniations. INM herniations containing PEVs and those lacking PEVs were scored together. INM herniations were more frequently observed in cells infected with viruses deficient in pUs3 than in cells infected with WT virus or pUL16 deficient virus. The number of nuclear sections scored were: WT 186 (n=12); ΔUs3 (n=28); Δ16 (n=14); Δ16/ΔUs3 isolate 1 (n=38) and Δ16/ΔUs3 isolate 2 (n=10). Error bars are standard error of the mean.

## Discussion

As pUL21 and pUL16 both function in the nuclear egress of HSV-2, this study was initiated to determine the effect of deleting these proteins on the localization of NEC components. Since pUL21 and pUL16 are capable of forming a complex (22), we anticipated that the NEC localization phenotype would be similar between the Δ21 and Δ16 strains. This was not the case (Fig 2). Whereas Δ21 demonstrated profound NEC mislocalization, the NEC localization in Δ16 infected cells was indistinguishable from that observed in WT infected cells. These findings suggested that pUL21 and pUL16 were functioning in mechanistically distinct ways to promote nuclear egress and that it was unlikely that a pUL21/pUL16 complex was the functional unit enabling efficient nuclear egress.

How might pUL16 function in HSV-2 nuclear egress? A failure to produce DNA containing capsids in pUL16 mutant infected cells cannot explain this defect (Fig 11C) (19, 20). It is equally unlikely that the lack of pUL16 impacts the transport of DNA-containing capsids to the INM as capsid movement within the nucleus is thought to occur primarily by diffusion (25, 26). It may be that the absence of pUL16 influences the composition of nuclear capsids such that they are unable to efficiently engage the NEC.

Analysis of cells infected with UL21 mutants derived from multiple HSV-2 and HSV-1 strains indicated that the impact of pUL21 loss on NEC localization was conserved between HSV strains and species. Ultrastructural examination of UL21 mutant strains revealed a number of nuclear envelope perturbations including extravagations of the nuclear envelope into the cytoplasm as well as invaginations of the nuclear envelope and the INM into the nucleoplasm (Fig 6). When INM invaginations were observed, PEVs were often seen accumulating in these structures that were reminiscent of INM herniations seen in cells infected with Us3 mutants derived from PRV (17), EHV-1 (15), MDV (16), HSV-1 (3) and HSV-2 (Fig 11A). Knockdown of pUL31 in cells infected with Δ21 prevented these nuclear envelope perturbations indicating that pUL31 activity was required to drive the formation of these structures and further suggesting that NEC activity is dysregulated in the absence of pUL21.

Co-expression of pUL31 and pUL34 in the absence of other viral proteins leads to vesiculation of the INM at the nuclear periphery leading to the formation of perinuclear vesicles (12), and the interaction of purified pUL31 and pUL34 with synthetic membranes also results in membrane vesiculation (8, 13). Evidence is mounting that the membrane vesiculation activity of the NEC is regulated in the context of viral infection (27). In cells infected with WT strains of HSV-1 (Fig 3 and (2)), PRV (28) and HSV-2 (Fig 3 and (29)), pUL31 and pUL34 localize in a smooth and even distribution at the nuclear membrane and INM vesiculation is rarely apparent, suggesting that factors found in virally-infected cells regulate NEC activity such that it is only active when capsids are presented for envelopment. One such regulator of the NEC is the viral serine/threonine kinase pUs3. Although pUs3 functions in multiple facets of the virus-host interaction (30), the best characterized is its role in nuclear egress. PEVs were observed to accumulate in INM herniations in cells infected with a PRV Us3 mutant virus (17). The interpretation of this observation was that PEV accumulations result as a consequence of the failure of PEVs to de-envelop at the ONM. This interpretation has been widely utilized to explain the phenotypes of Us3 mutant strains derived from other herpesviruses including HSV-1, EHV-1 and MDV (3, 15, 16). The data presented here confound this interpretation. Our analysis of two independently constructed Δ16/ΔUs3 mutants revealed the presence of INM herniations that lack PEVs (Fig 11C). Collectively, these findings suggest that formation of INM herniations is not driven by the accumulation of PEVs in the perinuclear space but rather by the absence of pUs3.

Baines and co-workers demonstrated that pUs3 phosphorylates multiple serine residues in the N-terminus of pUL31 and that serine to alanine substitution of these residues recapitulates the Us3 null phenotype (i.e. the accumulation of PEVs in the perinuclear space) (18). We hypothesize that the absence of pUs3-mediated phosphorylation of pUL31 “activates” the NEC to form INM herniations. We suggest that, by breaching the lamina and marginalized chromatin, INM herniations serve as attractive sites for primary envelopment. However, this might position PEVs too far away from the ONM where de-envelopment must take place and thereby leads to their accumulation in the perinuclear space. In other words, if the separation between the site of primary envelopment and the site of de-envelopment is too large, this can impair nuclear egress. This idea is supported by observations from the Mettenleiter laboratory showing that when the perinuclear space was enlarged through expression of a dominant-negative SUN protein, PRV PEVs accumulated within this compartment (31). As formation of INM invaginations is a feature associated with UL21 mutants (Fig 6), separation of the sites of primary envelopment and de-envelopment may also explain why UL21 mutants can display nuclear egress impairment even in the presence of functional pUs3.

Our observation that knockdown of pUL31 in Δ21 infected cells prevented NEC mislocalization implies that NEC activity is required for this mislocalization and further, similar to pUs3, pUL21 can regulate NEC activity. In addition to other nuclear envelope perturbations, we observed the accumulation of PEVs within extensive INM invaginations, particularly in HSV-1 strains harboring UL21 deletions. As mentioned above, a likely explanation for the Us3 mutant PEV accumulation phenotype is the hypo-phosphorylation of pUL31 observed in the absence of pUs3 (18). It may be that pUL21 enhances pUs3 activity or, alternatively, regulates the phosphorylation status of pUL31 by other means such as by preventing its dephosphorylation. Our earlier observations that pUL21 localizes to the nuclear rim during virus infection would position pUL21 in the appropriate location to perform such activities (21). Determining the mechanism by which pUL21 localizes to the nuclear rim and examination of pUL31 phosphorylation status in UL21 mutant infected cells should provide clearer insight into how pUL21 regulates NEC activity.

Our observation (Fig 2 and 11A) that the NEC was mislocalized and that PEVs accumulated in INM herniations in ΔUs3 infected cells lies in contrast with work from the Kawaguchi laboratory, which suggested that nuclear egress is unperturbed in HSV-2 Us3 mutants (29). As the strain of HSV-2 and the cell type utilized in the two laboratories were identical, the nature of the mutations may be responsible for this discrepancy. The ΔUs3 strain used herein contains tandem nonsense codons after the second Us3 codon and infections with ΔUs3 produce no pUs3 (Fig 10, Supplemental Fig 1). By contrast, the strains utilized by Morimoto and colleagues express catalytically inactive forms of pUs3 raising the possibility that pUs3 is important for HSV-2 nuclear egress, but its kinase activity is not (29). Interestingly, replacement of HSV-1 Us3 with HSV-2 Us3 resulted in a chimeric strain that displayed nuclear egress deficiencies (32). These findings indicated that HSV-2 Us3 could not complement the loss of HSV-1 Us3 and further supported the idea that HSV-2 pUs3 does not function in nuclear egress. However, based on our findings that the absence of pUs3 in HSV-2 perturbs nuclear egress (Figs 2 and Fig 11A, B), it may be that HSV-2 pUs3 expressed in the HSV-1 background strain is unable to engage other, as yet unidentified, viral or cellular proteins required for pUs3 nuclear egress activity.

In conclusion, we have provided evidence that pUL16, pUL21 and pUs3 play distinct roles in the nuclear egress of HSV-2 capsids. Whereas, pUL16 appears to function upstream of capsid primary envelopment, both pUs3 and pUL21 appear to function by regulating NEC activity through distinct mechanisms.

## Materials and methods

### Cells and viruses

African green monkey kidney cells (Vero), 293T cells, HeLa cells, human keratinocytes (HaCaT) and murine L fibroblasts were maintained in Dulbecco’s modified Eagle medium (DMEM) supplemented with 10% fetal bovine serum (FBS) in a 5% CO_2_ environment. L cells and HaCaT cells that stably produce pUL21 of HSV-2 strain 186 (referred to herein as L21 and HaCaT21) and L cells and HaCaT cells that stably produce pUL16 of HSV-2 strain 186 (referred to herein as L16 and HaCaT16) were isolated by retroviral transduction using an amphotropic Phoenix-Moloney murine leukemia virus system, as described previously (19, 21). HSV-2 strains SD90e, HG52, and 186 were acquired from Dr. D. M. Knipe (Harvard University), Dr. D. J. McGeoch (MRC Virology Unit, University of Glasgow), and Dr. Y. Kawaguchi (University of Tokyo), respectively; HSV-1 strains KOS and F were acquired from Dr. L. W. Enquist (Princeton University). Viruses deficient in pUL21 or pUL16 were constructed by two-step Red-mediated mutagenesis of a bacterial artificial chromosome (BAC) or by CRISPR/Cas9 mutagenesis as described previously (19–21, 23). HSV-2 186 virus deficient in pUs3 (ΔUs3) was constructed by two-step Red-mediated mutagenesis (33), using pYEbac373 (21) in *Escherichia coli* GS1783. Primers 5′-CCCGTCGCTCGGGGTGCTCGTTGGTTGGCACGCGCGACGCGGCGA<u>ATGGCC</u>**TGATGA**AAGAGGATGACGACGATAAGTAGGG-3′ and 5′-CTTGTCGGGTCTACGGTAGACCCCACAGAACTTTCATCAGGCCATTCGCCGCGTCGCGCGCAACCAATTAACCAATTCTGATTAG-3′ were used to amplify a PCR product from pEP-Kan-S2, a kind gift of Dr. N. Osterrieder (Freie Universität Berlin), to introduce two tandem stop codons (bold) after the first two codons of Us3 (underlined).

Restriction fragment length polymorphism analysis was used to confirm the integrity of each recombinant BAC clone in comparison to the wild type (WT) BAC by digestion with EcoRI. Additionally, a PCR fragment that spanned the region of interest was amplified and sequenced to confirm the presence of the tandem stop codons in Us3. Virus was reconstituted from ΔUs3 BAC DNA as described previously (21). To repair the Us3 deletion mutant, primers 5’-CCCGTCGCTCGGGGTGCTCGTTGGTTGGCACGCGCGACGCGGCGAATGGCCTGTCGTAAGAGGATGACGACGATAAGTAGGG-3′ and 5’-CTTGTCGGGTCTACGGTAGACCCCACAGAACTTACGACAGGCCATTCGCCGCGTCGCGCGCAACCAATTAACCAATTCTGATTAG-3′ were used to amplify a PCR product from pEP-Kan-S2, which was used to restore the complete Us3 gene in the ΔUs3 BAC as described above. Western blot analysis confirmed that pUs3 was not produced in cells infected with ΔUs3 (Supplemental Fig 1) and was restored in the repaired strain ΔUs3R. HSV-2 186 viruses deficient in both pUs3 and pUL16 (Δ16/ΔUs3) were constructed by CRISPR/Cas9 mutagenesis with HSV-2 UL16 specific guide RNAs on viral genomic DNA isolated from ΔUs3 as described previously (20). Two separate Δ16/ΔUs3 viruses, isolated from independent co-transfections, were utilized in this study. In both Δ16/ΔUs3 viruses, the UL16 gene contains an in-frame deletion of codons 10 through 360. WT viruses and pUs3 deficient virus were propagated in either Vero or HaCaT cells. All pUL21 deficient viruses were propagated in either L21 or HaCaT21 cells. All pUL16 deficient viruses were propagated in L16 cells or HaCaT16 cells. Times post infection, reported as hours post infection (hpi), refer to the time elapsed following medium replacement after a one hour inoculation period.

### Production of antisera against HSV-2 pUL31 and HSV-2 pUL34

Recombinant full length HSV-2 GST-UL31 and HSV-2 GST-UL34 fusion proteins were produced in *E. coli* strain Rosetta(DE3). Bacteria were lysed and inclusion bodies were purified using a B-Per protein purification kit (Thermo Scientific, Rockford, IL) according to the manufacturer’s instructions. Proteins in inclusion bodies were separated on preparative SDS-PAGE gels, the bands corresponding to GST fusions were excised and sent to Cedarlane (Burlington, ON) to immunize rats for pUL31 antiserum production or to Virusys (Taneytown, MD) to immunize chickens for pUL34 antiserum production. Rat polyclonal antiserum against pUL31 was used for indirect immunofluorescence microscopy at a dilution of 1:500 and for Western blotting at a dilution of 1:200; chicken polyclonal antiserum against pUL34 was used for indirect immunofluorescence microscopy at a dilution of 1:500 and for Western blotting at a dilution of 1:500.

### Other immunological reagents

Alexa Fluor 488 conjugated donkey anti-rat IgG, Alexa Fluor 568 goat anti-rat IgG, and Alexa Fluor 568 conjugated goat anti-chicken IgG (Molecular Probes, Eugene, OR) were all used for indirect immunofluorescence microscopy at a dilution of 1:500. Rabbit polyclonal antiserum against pUL16 (34), a kind gift from Dr. J.W. Wills (The Pennsylvania State University College of Medicine), was used for Western blotting at a dilution of 1:3,000; rat polyclonal antiserum against pUL21 (21) was used for Western blotting at a dilution of 1:600; rat polyclonal antiserum against pUs3 (35) was used for Western blotting at a dilution of 1:1,000; mouse monoclonal antibody against ICP27 (Virusys, Taneytown, MD) was used for Western blotting at a dilution of 1:500, mouse monoclonal antibody against EGFP (Clontech, Mountain View, CA) was used for Western blotting at a dilution of 1:1,000; mouse monoclonal antibody against ICP8 (Virusys, Taneytown, MD) was used for Western blotting at a dilution of 1:4,000 and mouse monoclonal antibody against β–actin (Sigma, St. Louis, MO) was used for Western blotting at a dilution of 1:2,000. Horseradish peroxidase-conjugated goat anti-mouse IgG, horseradish peroxidase-conjugated goat anti-chicken IgY, horseradish peroxidase-conjugated rabbit anti-rat IgG and horseradish peroxidase-conjugated goat anti-rabbit IgG (Sigma, St. Louis, MO) were used for Western blotting at dilutions of 1:10,000, 1:30,000, 1:80,000 and 1:5,000, respectively.

### Indirect immunofluorescence microscopy

Cells for microscopic analyses were grown on 35 mm glass bottom dishes (MatTek, Ashland, MA), fixed with 4% paraformaldehyde and stained as described previously (35). Images were captured with an Olympus FV1000 laser scanning confocal microscope using a 60X (1.42 NA) oil immersion objective and Fluoview 1.7.3.0 software. Composites of representative images were prepared using Adobe Photoshop software.

### Transmission electron microscopy (TEM)

Vero cells growing in 100-mm dishes were infected with virus at a multiplicity of infection (MOI) of 3 and processed for TEM at 16 hpi. Infected cells were rinsed with PBS three times before fixing in 1.5 ml of 2.5% EM grade glutaraldehyde (Ted Pella, Redding, CA) in 0.1 M sodium cacodylate buffer (pH 7.4) for 60 minutes. Cells were collected by scraping into fixative and centrifugation at 300Xg for 5 minutes. Cell pellets were carefully enrobed in an equal volume of molten 5% low-melting temperature agarose (Lonza, Rockland, ME) and allowed to cool. Specimens in agarose were incubated in 2.5% glutaraldehyde in 0.1 M sodium cacodylate buffer (pH 7.4) for 1.5 hours and post-fixed in 1% osmium tetroxide for 1 hour. The fixed cells in agarose were rinsed with distilled water 3 times and stained in 0.5% uranyl acetate overnight before dehydration in ascending grades of ethanol (30%-100%). Samples were transitioned from ethanol to infiltration with propylene oxide and embedded in Embed-812 hard resin (Electron Microscopy Sciences, Hatfield, PA). Blocks were sectioned at 50-60 ηm and stained with uranyl acetate and Reynolds’ lead citrate. Images were collected using a Hitachi H-7000 transmission electron microscope.

### DsiRNA knockdown

Dicer-substrate siRNAs (DsiRNAs) directed against HSV-2 strain 186 *UL31* were custom designed using the Integrated DNA Technologies (IDT) design tool (IDT, Coralville, IA). DsiRNAs were transfected into Vero cells growing on 35 mm glass bottom dishes or on standard 35 mm dishes using PepMute transfection reagent (SignaGen Laboratories, Rockville, MD) according to manufacturer’s protocols. Twenty-four hours after transfection, transfected cells were infected with virus at a MOI of 0.5. At 18 hpi, infected cells were fixed in preparation for indirect immunofluorescent staining or whole cell lysates were prepared for Western blot analysis.

### Plasmids and transfections

The construction of a plasmid encoding EGFP-UL31 was described previously (36). To construct EGFP-UL34, PCR utilizing a forward primer 5’-AGTTCGAATTCTATGGCGGGGATGGGGAAGCCCTACG-3’ and reverse primer 5’-GATCGTCGACTCATATAGGCGCGCGCCAACCGCC-3’ were used to amplify the UL34 gene using purified HSV-2 DNA as template. The product was digested with EcoRI and SalI and ligated into similarly digested pEGFP-C1 (Clontech Laboratories, Mountain View, CA). All plasmids constructed utilizing PCR were sequenced to ensure that no unintended mutations were introduced. For preparing whole cell extracts of transfected cells for Western blot analysis, plasmids were transfected into 293T cells using the calcium phosphate co-precipitation method (37). To examine the localization of EGFP fusion proteins by microscopy, plasmids were transfected into HeLa cells growing on 35 mm glass bottom dishes using X-treme GENE HP DNA transfection reagent (Roche, Laval, QC) according to manufacturer’s protocols.

### Preparation and analysis of whole cell protein extracts

To prepare whole cell protein extracts of transfected or infected cells for Western blot analyses, cells were washed with cold phosphate buffered saline (PBS) then scraped into cold PBS containing protease inhibitors (Roche, Laval, QC) and 5 mM sodium fluoride (New England Biolabs, Ipswich, MA) and 1 mM sodium orthovanadate (New England Biolabs, Ipswich, MA) to inhibit phosphatases. Harvested cells were transferred to a 1.5 ml microfuge tube containing 3X SDS-PAGE loading buffer. The lysate was repeatedly passed through a 28 1/2 gauge needle to reduce viscosity and then heated at 100°C for 5 min. For Western blot analysis, 10 to 20 μl of whole cell extract was electrophoresed through SDS-PAGE gels. Separated proteins were transferred to PVDF membranes (Millipore, Billerica, MA) and probed with appropriate dilutions of primary antibody followed by appropriate dilutions of horseradish peroxidase conjugated secondary antibody. The membranes were treated with Pierce ECL Western Blotting Substrate (Thermo Scientific, Rockford, IL) and exposed to film.

## Acknowledgments

We thank Dr. Xiaohu Yan (Queen’s University) for assistance with preparing samples for TEM analyses and Kristin Piche for technical assistance with plasmid construction.

---

**Supplemental Fig 1.**
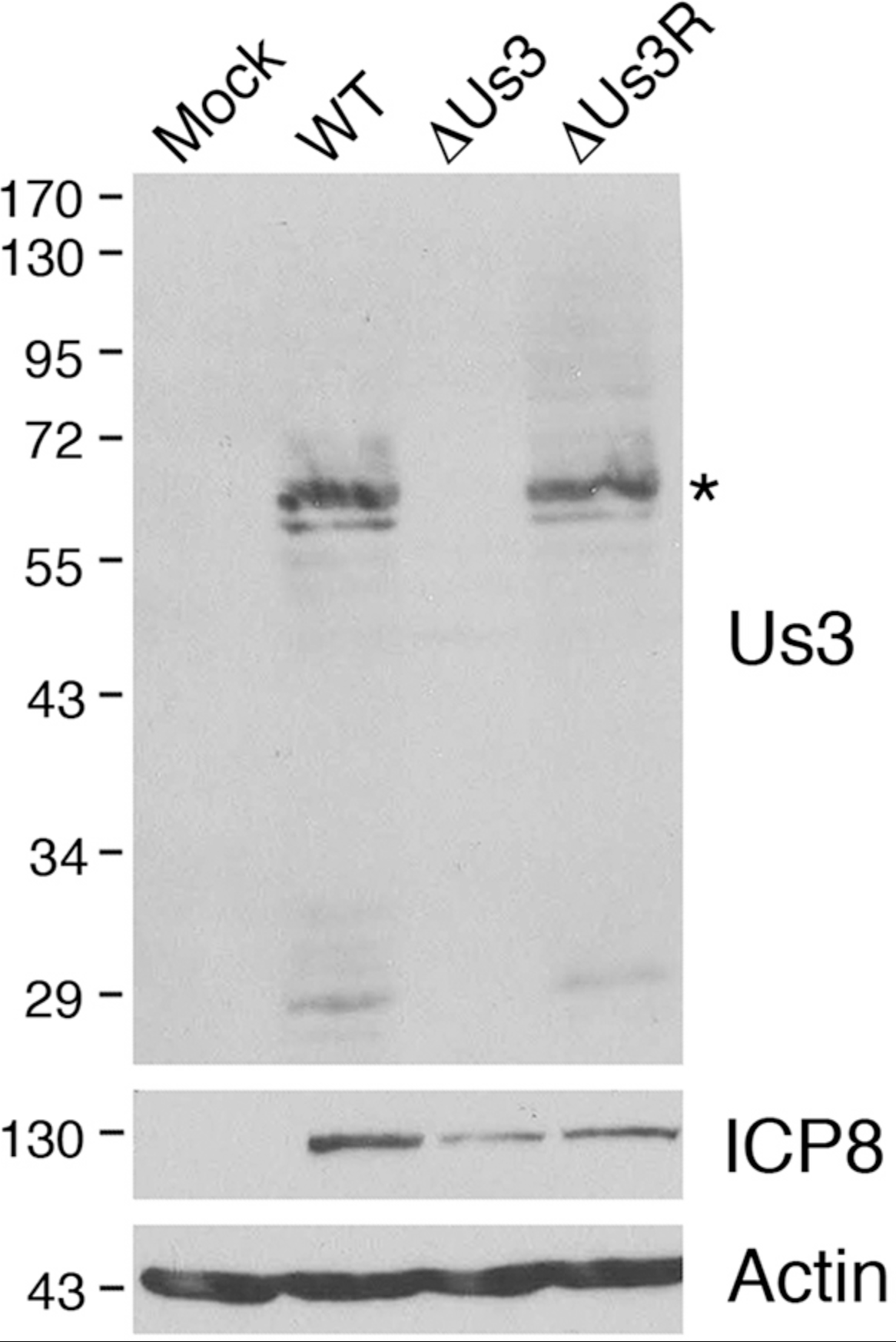
Characterization of ΔUs3 and ΔUs3R strains. Vero cells were infected with the indicated viruses at an MOI of 0.1. At 18 hpi, whole cell extracts were prepared and electrophoresed through 10% polyacrylamide gels and transferred to PVDF membranes. Membranes were probed with antisera indicated on the right side of each panel. Molecular weight markers in kDa are indicated on the left side of each panel. Asterisk indicates the position of full length pUs3.

